# A High Content Screen for Mucin-1-Reducing Compounds Identifies Fostamatinib as a Candidate for Rapid Repurposing for Acute Lung Injury during the COVID-19 pandemic

**DOI:** 10.1101/2020.06.30.180380

**Authors:** Maria Alimova, Eriene-Heidi Sidhom, Abhigyan Satyam, Moran Dvela-Levitt, Michelle Melanson, Brian T. Chamberlain, Seth L. Alper, Jean Santos, Juan Gutierrez, Ayshwarya Subramanian, Elizabeth Grinkevich, Estefania Reyes Bricio, Choah Kim, Abbe Clark, Andrew Watts, Rebecca Thompson, Jamie Marshall, Juan Lorenzo Pablo, Juliana Coraor, Julie Roignot, Katherine A. Vernon, Keith Keller, Alissa Campbell, Maheswarareddy Emani, Matthew Racette, Silvana Bazua-Valenti, Valeria Padovano, Astrid Weins, Stephen P. McAdoo, Frederick W.K. Tam, Lucienne Ronco, Florence Wagner, George C. Tsokos, Jillian L. Shaw, Anna Greka

## Abstract

Drug repurposing is the only method capable of delivering treatments on the shortened time-scale required for patients afflicted with lung disease arising from SARS-CoV-2 infection. Mucin-1 (MUC1), a membrane-bound molecule expressed on the apical surfaces of most mucosal epithelial cells, is a biochemical marker whose elevated levels predict the development of acute lung injury (ALI) and respiratory distress syndrome (ARDS), and correlate with poor clinical outcomes. In response to the pandemic spread of SARS-CoV-2, we took advantage of a high content screen of 3,713 compounds at different stages of clinical development to identify FDA-approved compounds that reduce MUC1 protein abundance. Our screen identified Fostamatinib (R788), an inhibitor of spleen tyrosine kinase (SYK) approved for the treatment of chronic immune thrombocytopenia, as a repurposing candidate for the treatment of ALI. *In vivo*, Fostamatinib reduced MUC1 abundance in lung epithelial cells in a mouse model of ALI. *In vitro*, SYK inhibition by Fostamatinib promoted MUC1 removal from the cell surface. Our work reveals Fostamatinib as a repurposing drug candidate for ALI and provides the rationale for rapidly standing up clinical trials to test Fostamatinib efficacy in patients with COVID-19 lung injury.

## Introduction

Drug repurposing is a strategy to identify novel uses for approved or investigational drugs outside the scope of their originally designated purposes. This approach offers several advantages over *de novo* drug development (Ashburn and Thor, 2004; Pushpakom et al., 2019). First and foremost, the risk of toxicity is much lower as repurposed approved drugs have been proven safe for human use in the original indication. Second, and of critical importance for addressing the global public health crisis attributed to SARS-CoV-2, is that drug repurposing offers the only method for delivering treatments on the shortened time-scale required to treat COVID-19 patients. Current management of COVID-19 is largely supportive and with severely limited therapeutic options. Once infection with SARS-CoV-2 is established, a subset of patients experience severe complications such as acute respiratory distress syndrome (ARDS), an extreme form of acute lung injury characterized by disruption to the alveolar epithelium (Ruan et al., 2020; Zhou et al., 2020). ARDS is a life-threatening condition with mortality rates as high as 40% (Acute Respiratory Distress Syndrome et al., 2000; Determann et al., 2010; Rubenfeld et al., 2005). COVID-19-associated ARDS is often fatal, especially in the presence of several pre-existing conditions. The currently limited therapeutic interventions available for COVID-19 (Cao et al., 2020; Grein, 2020) have contributed to an estimated 400,000 deaths worldwide at the time of writing (Dong et al., 2020). Identification of drugs with efficacy in treating ALI in severely affected COVID-19 patients remains an urgent need.

ARDS patients exhibit high serum levels of mucin-1/MUC1 (KL-6) (Nakashima et al., 2008). MUC1 is a transmembrane protein expressed on the apical membrane of most mucosal epithelial cells and plays a critical role in lining the airway lumen (Kato et al., 2017). Mucins are glycoproteins that impart specific properties to mucus. In response to specific stimuli, goblet cells can rapidly secrete mucus by exocytosis to form a mucus layer that lines the airways. In healthy individuals, mucus along the lumen serves as a major protective barrier against inhaled pathogens, toxins, and other foreign particles. However, excessive mucus in the airways has been linked to increased frequency and duration of infections, decreased lung function, and increased mortality from respiratory diseases (Vestbo, 2002). Abnormalities in mucus production contribute to severe pulmonary complications and death from respiratory failure in patients with diseases such as cystic fibrosis, chronic obstructive pulmonary disease (COPD), and acute lung injury due to viral pathogens, such as SARS-CoV2. Elevated serum KL-6/MUC1 levels are an early prognostic marker of the therapeutic effect of high-dose corticosteroids in patients with rapidly progressing idiopathic pulmonary fibrosis (Yokoyama et al., 1998). Serum KL-6/MUC1 levels are also elevated in patients with interstitial pneumonitis (Ishikawa et al., 2012; Kohno, 1999). Moreover, transgenic mice expressing human MUC1 and subjected to LPS-induced ALI exhibit elevated KL-6 both in alveolar pneumocytes and in serum (Sakai et al., 2013).

Prompted by the connection between elevated MUC1 and ALI, we investigated the possibility of identifying MUC1-reducing drugs for rapid repurposing. We had originally screened the Broad Repurposing Library (comprised of 3,713 compounds at different stages of pre-clinical and clinical development (Corsello et al., 2017)) to identify compounds capable of reducing a mutant MUC1 neo-protein (MUC1-fs) causing autosomal dominant tubulo-interstitial kidney disease-*mucin1* (ADTKD-*MUC1* or *MUC1* kidney disease, MKD) (Dvela-Levitt et al., 2019). In this context wildtype MUC1 (MUC1-wt) served as a control, as we sought compounds that specifically reduced the mutant, but not the wildtype form of MUC1. As the number of COVID-19 cases increased globally, we turned our attention to identifying MUC1-reducing compounds and mined this dataset to identify approved drugs that reduce expression of MUC1-wt. We searched for MUC1-reducing compounds based on the following criteria: 1) a drug that reduces MUC1-wt protein in a dose-dependent manner; 2) a drug with a favorable toxicity profile; 3) a drug that reduces MUC1-wt by non-transcriptional mechanisms (Dvela-Levitt et al., 2019), unlike transcriptional suppressors such as vitamin D agonists that have proven ineffective in the clinic (Castro et al., 2014); and 4) a drug that is US Food and Drug Administration (FDA)-approved. Based on these criteria, our screen identified R406, the active metabolite of Fostamatinib (R788, an oral prodrug rapidly converted to R406), as a repurposing candidate for the treatment of ALI.

## Results

### The FDA approved SYK inhibitor R406 depletes MUC1 from epithelial cells without affecting cell viability

To investigate the expression pattern of MUC1 in human tissue, we took advantage of the openly available Human Protein Atlas (HPA)(www.proteinatlas.org) (Uhlen et al., 2015). Immunoperoxidase staining of human lung showed MUC1 expression in alveolar epithelium. This finding confirmed multiple reports of MUC1 expression in normal and diseased human lung (Figure 1A) (Ishizaka et al., 2004; Ohtsuki et al., 2007). These data were corroborated by the expression of MUC1 mRNA in human lung reported by the HPA, GTEx and FANTOM5 databases (Figure 1B) and by the Human Lung Atlas project (Muus, 2020)

**Figure 1.**
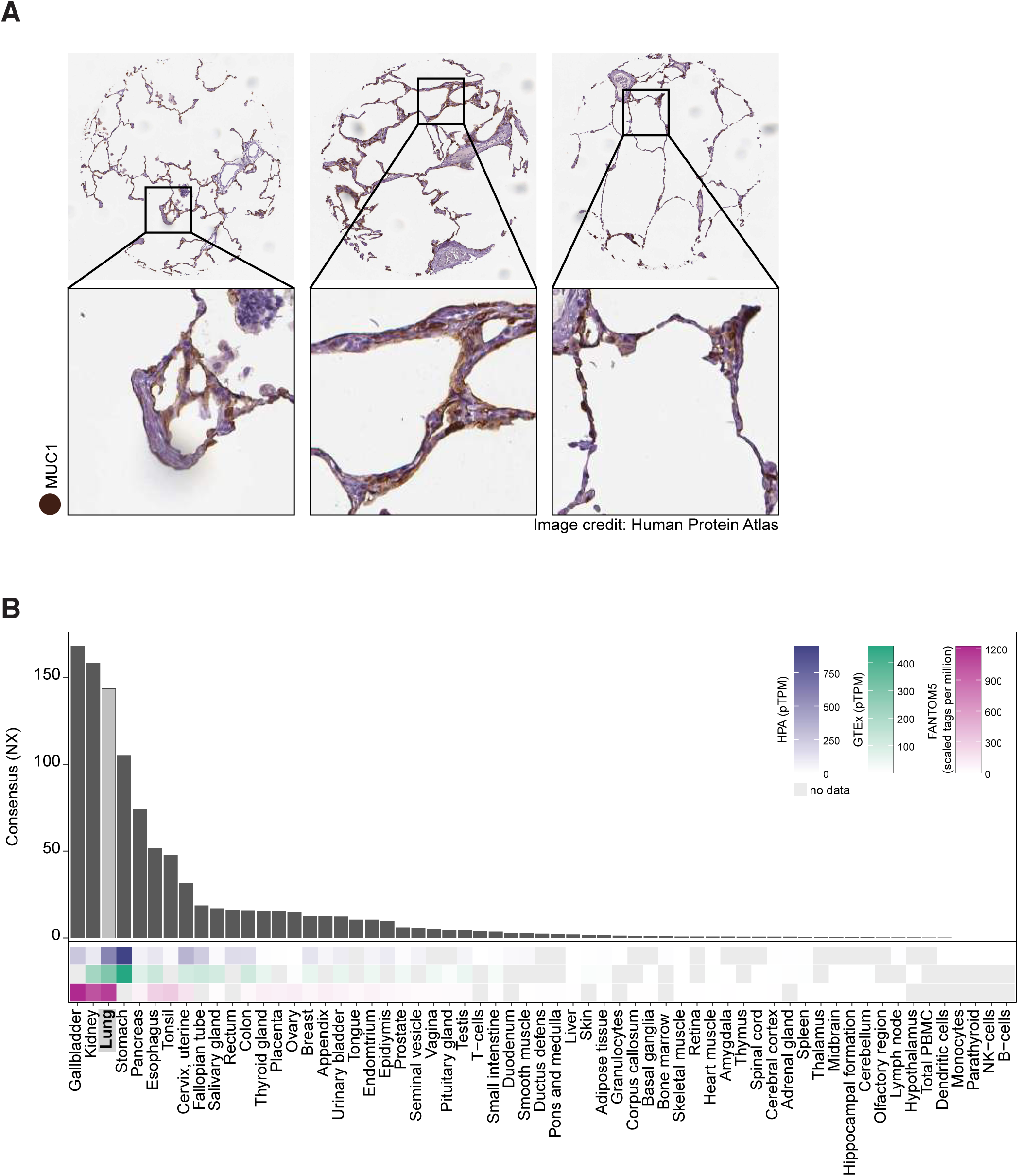
High relative expression of MUC1 in human lung. A. Immunoperoxidase staining of three human lung samples demonstrates MUC1 protein abundance in lung tissue. B. mRNA expression data from three datasets (HPA, GTEx, and FANTOM5, heatmaps) combined into a CONSENSUS normalized transcript expression level (bar plot) show enhanced expression of MUC1 mRNA in human lung. NX, normalized expression; pTPM, protein-coding transcripts per million.

We screened 3,713 compounds of the Repurposing Library for their ability to reduce MUC1 protein levels (Figure 2A). The screen employed high-content immunofluorescence (IF) imaging of an immortalized kidney tubular epithelial cell line (P cells) that express endogenous MUC1 on the plasma membrane, to simultaneously assess MUC1 protein abundance and cell number as an index of cell toxicity (Dvela-Levitt et al., 2019). The bromodomain inhibitor JQ1 served as positive control, as preliminary experiments demonstrated complete transcriptional suppression of MUC1 by JQ1. Each compound in the Repurposing Library was tested in a 5 concentration, 10-fold dilution series with a top concentration of 35 µM in the initial screen. Positive hits from the 5-dose screen were defined by two criteria: lack of cellular toxicity (less than 20% reduction in cell number compared to the DMSO control); and reduction of MUC1 abundance by >30% (normalized to DMSO and JQ1 controls) at 2 or more consecutive non-toxic concentrations of test compound. The compounds that met these criteria included two major groups, bromodomain inhibitors (blue) and vitamin D receptor agonists (orange, Figure 2B). Our screen also identified drugs that increased MUC1 levels (Figure 2B), including glucocorticoid receptor agonists. Most compounds that increased MUC1, such as epidermal growth factor receptor (EGFR) inhibitors (green), also reduced cell number, indicating an association between cell toxicity and increased MUC1 levels (Figure 2B).

**Figure 2.**
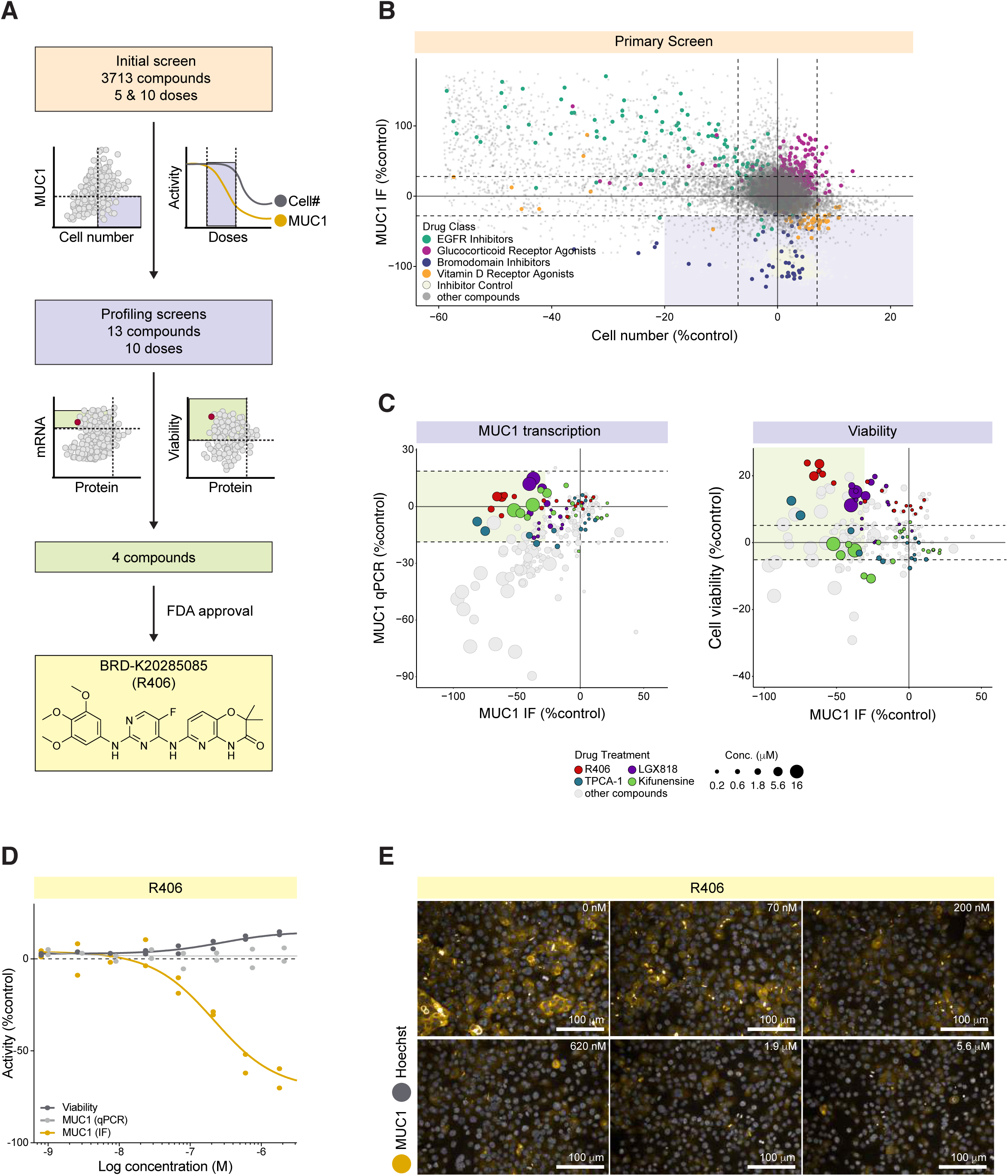
High Content Screening reveals significant and dose-dependent reduction in MUC1 by the FDA-approved SYK inhibitor R406. A. Screening pipeline. B. Primary screen revealed four major groups of compounds which affected MUC1 levels. MUC1 immunofluorescence (IF) signal intensity per cell (normalized to positive control JQ1 minus DMSO-treated controls) plotted vs. DMSO-normalized cell number. Horizontal and vertical dashed lines delineate mean DMSO values +/-3*SD for both MUC1 intensity and cell number. Lavender-shaded area demarcates candidate MUC1 suppressors. C. qPCR and cell viability profiling screens identified four compounds that reduced MUC1 protein abundance without changing MUC1 mRNA level, and in the absence of cytotoxicity. Left: MUC1 signal intensity per cell plotted vs. MUC1 mRNA level (qPCR assay). Both parameters are normalized to JQ1 minus DMSO-treated controls. Right: JQ1 minus DMSO-normalized MUC1 signal intensity per cell plotted vs. DMSO-normalized cell viability (a number of viable cells after 6 days exposure to the test compounds). Horizontal dashed lines delineate SD for DMSO treated control wells for both cell viability and MUC1 qPCR. Green-shaded areas demarcate candidate hits. D. R406 concentration response curves for MUC1 protein abundance (black), MUC1 mRNA abundance (light gray), and cell viability (dark gray). E. MUC1 IF in kidney epithelial cells treated for 48 hours in the absence (DMSO) and presence of R406 at the indicated concentrations.

Two hundred and three compounds were re-tested at 10 concentrations to generate more complete dose response curves for each compound. In this screen, any compound that reduced MUC1 by >30% at 2 or more consecutive concentrations without evidence of toxicity (cell numbers within 20% of the DMSO control) was considered a positive hit. Thirteen hits from this screen were analyzed further in secondary profiling assays, including quantitative PCR (qPCR) and cell viability screens. MUC1 qPCR (Figure 2C) showed that for most compounds, reduction of MUC1 protein abundance (MUC1 IF) was highly correlated with parallel reductions in mRNA abundance (MUC1 qPCR). Of interest, 4 compounds reduced MUC1 protein without affecting MUC1 mRNA. These included the SYK inhibitor R406 (red, Figure 2C); the RAF inhibitor LGX818 (violet, Figure 2C); the mannosidase inhibitor Kifunensine (green, Figure 2C), and the IKK inhibitor TPCA-1 (turquoise, Figure 2C). None of these four compounds exhibited detectable toxicity (cell death or apoptosis) at any of the 10 tested concentrations. Importantly, only the SYK inhibitor R406 is FDA-approved (Figure 2A and D). No additional compounds with known activity against SYK met our screening criteria. R406 decreased MUC1 protein abundance in cells with an EC50 of approximately 200 nM. The lack of effect on MUC1 mRNA levels (as shown by qPCR) indicated that MUC1 protein reduction was not achieved via transcriptional repression (representative images of cells treated with R406 at a range of concentrations; Figure 2E). The parent molecule of R406, Fostamatinib (R788), showed no activity in the initial 5-concnetration screen.

### R406 preferentially depletes MUC1 from the plasma membrane

Image analysis from the high content screen revealed that, while MUC1 was preferentially localized to the plasma membrane in DMSO-treated control cells, treatment with 200 nM of R406 reduced MUC1 from the plasma membrane and redistributed a fraction of the protein to the intracellular compartment with a perinuclear distribution pattern (Figure 3A). We quantified this relocalization using a STAR morphology “Profile” Module (Star Methods) that allowed selective measurement of MUC1 signal distribution in cellular sub-compartments. Figure 3B shows an analysis sequence for the sub-compartment corresponding to the cell region closest to the plasma membrane. Within each cell, the nucleus border was identified based on Hoechst staining (blue, second panel, Figure 3B) and the plasma membrane (orange, second panel, Figure 3B) was identified based on the MUC1 signal. The calculated STAR morphology “Membrane Profile” image was generated as illustrated in Figure 3B, panel 3 (STAR Methods). The calculated STAR Membrane Profile values for MUC1 in each well were compared with total cell MUC1 intensity (Figure 3C). As shown by a local regression, while most compounds that decreased MUC1 did not affect MUC1 membrane localization, R406 produced preferential depletion of MUC1 from the plasma membrane region at all active concentrations.

**Figure 3.**
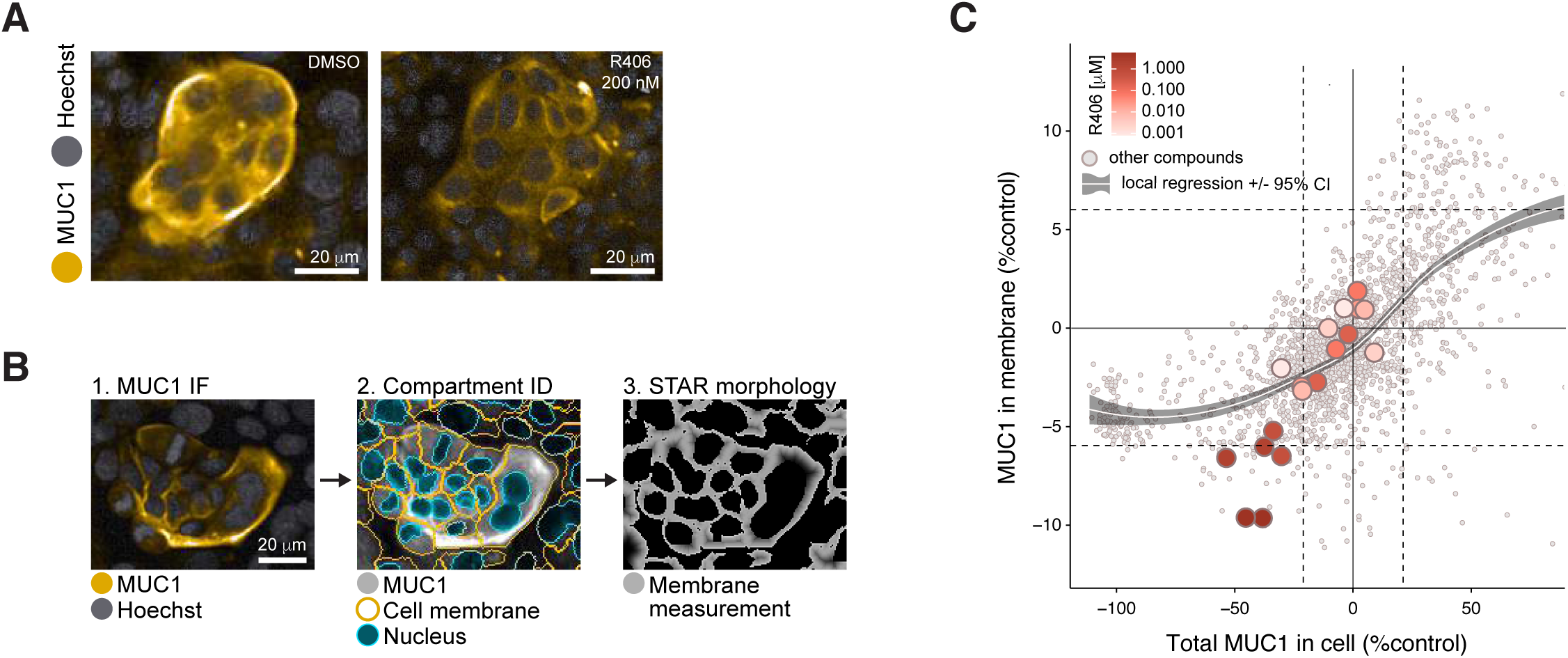
R406 preferentially depletes MUC1 from the plasma membrane. A. R406 (at EC50 concentration) substantially reduced MUC1 abundance in or near the plasma membrane, with a portion of MUC1 retained in cytosolic and perinuclear cell compartments. B. Image analysis for cell compartmentation using STAR morphology “Membrane - Profile” calculation (see Star Methods). Image 1: cells with MUC1 preferentially localized at plasma membrane; Image 2: Harmony software identification of nucleus (blue) and plasma membrane (gold) in each cell; Image 3: STAR morphology “Membrane Profile” for the MUC1 predominant localization within membrane compartment C. STAR morphology” Membrane Profile” analysis of 203 compounds screened at 10 doses. R406 at most active concentrations reduced plasma membrane MUC1 abundance to a greater degree than most other compounds, as shown by deviation from the local regression.□ MUC1 IF signal intensity per cell (normalized to JQ1 minus DMSO-treated controls) plotted vs. DMSO-normalized MUC1 predominance in plasma membrane region as calculated using the STAR morphology “Membrane Profile” module. Horizontal and vertical dashed lines delineate mean DMSO values +/- 2*SD for both plotted parameters. Local regression was calculated by locally estimated scatterplot smoothing (loess) method +/- 95% confidence interval (gray-shaded).

### *In vivo*, R406 reduces lung epithelial MUC1 in mice with ALI

SYK inhibition has previously been shown to suppress both local and remote lung injury (Pamuk, et al., 2010). R788 (fostamatinib disodium) is a methylene phosphate prodrug of R406 suitable for oral administration (McAdoo and Tam, 2011). To investigate whether administration of R788 might ameliorate ischemia-reperfusion (I/R)-induced remote lung injury by reducing MUC1 levels from the plasma membrane in the lung epithelium, C57BL/6J mice were fed a chow diet containing 3 grams/kilogram of R788 for 10 days. Immunohistochemical images obtained from formalin-fixed paraffin sections of lung tissues stained with MUC1, Phalloidin, and DAPI demonstrated that I/R-mediated ALI induced increased levels of MUC1 in lung epithelium, consistent with previous reports that excess MUC1 is injurious (Sakai et al., 2013). Importantly, MUC1 was significantly reduced by treatment with R788 (Figure 4A). Quantitative image analysis confirmed the *in vivo* efficacy of R788 in reducing MUC1 in injured lung epithelium (Figure 4B and C).

**Figure 4.**
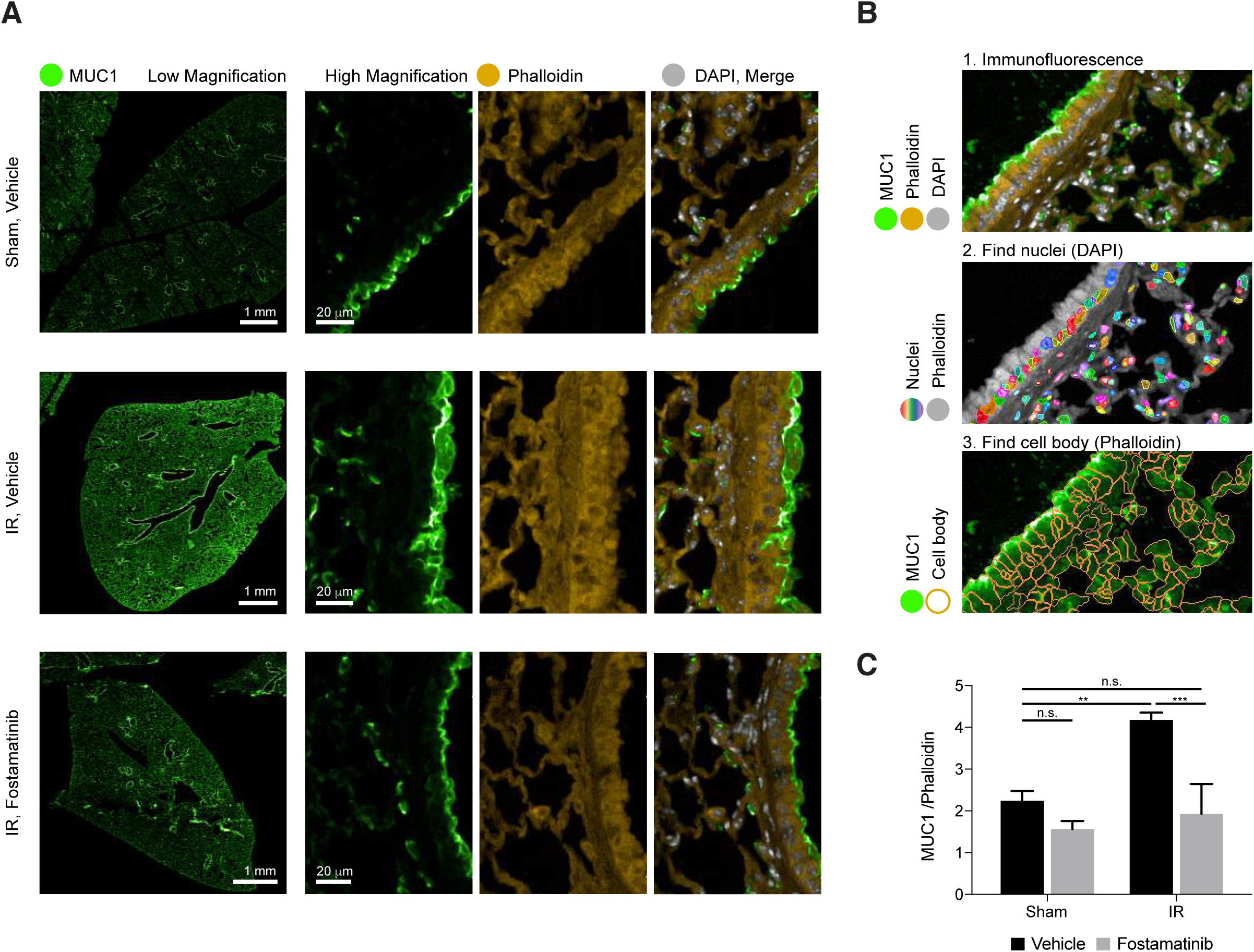
*In vivo*, R788 reduces excess MUC1 from lung epithelia of mice with ALI. A. Immunofluorescence images from lung tissue sections stained with MUC1 (green), Phalloidin (yellow), and DAPI (grey) demonstrate that ischemia-reperfusion (I/R)-induced remote ALI resulted in increased MUC1 in lung epithelium. Treatment with Fostamatinib over the course of 10 days suppressed MUC1 levels in mouse lung epithelium. B. Single cell tissue analysis, based on immunofluorescence of MUC1 and phalloidin (panel 1). In each image (panel 1) nuclei were identified based on DAPI staining (rainbow colors represent different cell nuclei in panel 2). The cell bodies were identified based on phalloidin staining surrounding each nucleus (orange cell borders, panel 3). Lastly, MUC1 IF intensity (green on panel 3) and Phalloidin intensities were calculated within each cell body. C. Bar graph ratio of MUC1/Phalloidin intensities in all cells of tissue sections from sham-treated mice and mice subjected to I/R-induced ALI, treated either with or without Fostamatinib. Average MUC1 intensity values per cell were normalized to the average phalloidin levels. Mean ± SD (n = 3 mice/condition/dose). ns, *p < 0.05 **p < 0.01 ***p < 0.001.

## Discussion

Our high-content screen identified R406, the active metabolite of Fostamatinib, as an FDA-approved candidate repurposing compound for the reduction of MUC1 protein levels in lung epithelium in the setting of ALI. R406 is a potent inhibitor of spleen tyrosine kinase (SYK), a cytosolic protein tyrosine kinase required for the expression of several proinflammatory cytokines (Yi et al., 2014). SYK is expressed in most leukocyte populations with roles in mediating signaling via classical immunoreceptors such as B-cell receptors and Fc receptors (Mocsai et al., 2010). SYK plays diverse roles in cellular adhesion, innate immune recognition, and platelet activation, and its central role in immune cell responses has made it a compelling target for the development of therapeutic agents. Over 70 patent filings describe small molecule inhibitors of SYK developed for treatment of diseases ranging from arthritis to asthma (Geahlen, 2014).

Fostamatinib is an effective treatment in experimental animal models of severe inflammatory diseases, including immune glomerulonephritis (McAdoo et al., 2014; Smith et al., 2010) and vasculitis (McAdoo et al., 2020). Phase II clinical trial results are expected assessing the effect of SYK inhibition in proliferative IgA nephropathy, an inflammatory kidney disease (NCT02112838). Fostamatinib was approved in April 2018 by the US Food and Drug Administration (FDA) for the treatment of chronic immune thrombocytopenia (ITP), an autoimmune disease that results in low levels of circulating platelets (Argade et al., 2015; Hilgendorf et al., 2011; Newland et al., 2018; Singh et al., 2012). Fostamatinib has also been extensively studied and found to be safe in more than 3000 patients with rheumatoid arthritis (Kunwar et al., 2016). In a Phase III clinical trial (Bussel et al., 2018), Fostamatinib was well tolerated at an oral dose of 100 mg twice daily. Mild or moderate adverse effects included diarrhea, hypertension, nausea, and an increase in alanine aminotransferase (ALT). These resolved spontaneously or with medical management, including antihypertensive or antimotility agents. This well-characterized clinical safety profile makes Fostamatinib an ideal candidate for rapid repurposing (Weinblatt et al., 2010).

Our finding that R406 preferentially depletes MUC1 abundance in or near the plasma membrane is consistent with a previously described mechanism by which SYK inhibition results in dephosphorylation of integral membrane proteins followed by their endocytic removal from the plasma membrane. For example, SYK signaling modulates CFTR abundance in human airway epithelial cell plasma membrane (Mendes et al., 2011). Interestingly, SYK in mucin-producing human NCI-H292 cells and in primary human nasal epithelial cells also regulates MUC5AC, a gel-like mucin that promotes lung epithelial injury (Na et al., 2016). Finally, in further support of the notion that MUC1 reduction is beneficial to injured lung epithelium, *Muc1* knockout in rat airway epithelial cells and MUC1 reduction in human lung epithelium resulted in diminished mucin hypersecretion and protection from lung injury (Kato et al., 2020).

Severe COVID-19 symptoms include viral-induced pneumonitis accompanied by prolonged, systemic cytokine release (Moore and June, 2020; Zhang et al., 2020) in which levels of interleukin-6 (IL-6) levels and other cytokines and acute phase reactants correlate with respiratory failure. Macrophage-derived IL-6 upregulates MUC1 in the human colon cancer HT-29 cell line (Li et al., 2009), suggesting a similar IL6-mediated upregulation of MUC1 may occur in SARS-CoV-2 infected lungs. A recent comparison of 15 hospitalized COVID-19 patients, 9 of whom were critically ill, to 28 critically ill patients with ARDS or sepsis found no statistically significant difference in circulating levels of IL-1b, IL-1RA, IL-6, IL-8 and TNF-alpha among these conditions (Wilson, 2020). These results indicate that COVID19-related ARDS is associated with inflammatory cytokine levels no higher than in ARDS due to other critical illnesses (Wilson, 2020). Another recent study analyzing serum concentrations of KL-6/MUC1 levels in hospitalized COVID19 patients suggested KL-6/MUC1 as a good prognostic biomarker of disease severity in COVID-19 patients (d’Alessandro et al., 2020). Given the roles of excess KL6/MUC1 in ALI and ARDS, we propose that Fostamatinib may confer benefit in patients with COVID-19 lung injury. In conclusion, our *in vitro* and *in vivo* data support the efficacy of Fostamatinib for the treatment of ALI. Here we provide a rationale for imminently designing and executing clinical trials to test whether repurposing this FDA-approved drug might confer clinical benefit for COVID-19 patients in the clinic.

## Acknowledgments

We gratefully acknowledge stimulating discussions with Broad colleagues Deborah Hung, Ramnik Xavier, Jay Rajagopal and Aviv Regev. We thank Terry Woo (BWH Pathology) and Stephen Straub (Perkin Elmer) for excellent technical support and expertise. SYK inhibitor R788 was provided as a gift to GCT by Rigel Pharmaceuticals (South San Francisco, CA).

This work was supported by PHS NIH R01AI148161 (GCT) and by the Slim Initiative for Genomic Medicine in the Americas (SIGMA), a collaboration of the Broad Institute with the Carlos Slim Foundation. FWKT is supported by the Ken and Mary Minton Chair of Renal Medicine.

## Author Contributions

MA, ASat, MDL, MM, JS, JG performed experiments; MA, EHS, ASat, MDL, BC, ASub performed data analysis and provided data visualization; SMcA, FT, GCT provided tissue and expertise in rodent models of lung disease; MA, EHS, MDL, BC, JS and AG wrote the manuscript; all authors participated in discussions on the scientific rationale for this study, read and approved the manuscript; AG supervised the project.

## STAR Methods

### Broad Repurposing Library

Quality control of The Broad Repurposing Library is performed at the time of plating by LCMS analysis using a Waters Acquity LC System with UV PDA detector and single quad (SQ) mass spectrometer R406 was confirmed present (MS (ESI/SQ) m/z: [M + H]+ Calculated for C22H23FN6O5+H: 471.2; Found: 470.9) in > 97% purity (PDA integration).

### Human Protein Atlas

MUC1 immunoperoxidase images were obtained from the Human Protein Atlas with the original source available at the following link: (https://www.proteinatlas.org/ENSG00000185499-MUC1/tissue/lung). mRNA expression data (https://www.proteinatlas.org/ENSG00000185499-MUC1/tissue) were downloaded from the Human Protein Atlas: (http://www.proteinatlas.org/ENSG00000185499.xml) (Uhlen et al., 2015). All graphs were visualized using ggplot2.

### Experimental Models

#### Cell Lines

Human P kidney epithelial cells (female) were previously generated from a patient with MUC1 kidney disease (Dvela-Levitt et al., 2019). The cells were maintained at 37°C with 5% CO2 in RenaLife Renal Basal Medium supplemented with RenaLife LifeFactors® (Lifeline Cell Technology), with the exclusion of Gentamycin and Amphotericin B. For all experiments, P cells were maintained below passage 12. The cells were generated with informed consent under WFUHS IRB00014033.

### Fluorescence image acquisition and analysis

All fluorescence imaging performed in this study was done using the Opera Phenix High-Content Screening System (PerkinElmer). For fluorescence imaging of cells (live cell or fixed cell imaging), CellCarrier 384-well Ultra microplates (Perkin Elmer) were used, and a minimum of nine fields was acquired per well using 20x water immersion objectives in a confocal mode.

Image analysis for all imaging experiments was performed using the Harmony software (PerkinElmer). Cell nuclei were first identified using Hoechst staining, and cell number was calculated. Cytoplasmic regions were then detected around each nucleus based on MUC1 channel. The cells from the edge of the field were eliminated from the analysis. For the quantification of MUC1 abundance, the total signal intensity value was calculated in the cell cytoplasm and the average signal per cell was calculated for each well.

For live cell image analysis, caspase 3/7 activation and/or DRAQ7 signal were used to detect cells going through apoptosis and/or cell death, respectively. Single cells were first identified using the digital phase contrast channel and cell number was calculated. Fluorescence intensities were then measured and the threshold for caspase 3/7 and DRAQ7 positive signal was determined. As an output, the number of live (neither caspase3/7 nor DRAQ7 signal detected) cells was calculated in each well at a particular time point.

For the MUC1 membrane prevalence, the images acquired during 10-dose screening were analyzed using the Harmony software STAR morphology feature, which calculates the signal distribution across different cell compartments. The inner side of plasma membrane compartment was analyzed by generating a “Membrane Profile Image” (profile 1/5 in Harmony software) (fig 3B). This function measures the closest distance of a given pixel to a cell border within a width of 4 pixels to preferentially weigh and quantify signal intensity (MUC1) closest to the plasma membrane signal.

For in vivo lung imaging, 20x water immersion objective was used, with 5% overlap for entire lung sections. Tissue Image Region for every tissue section was identified based on Gaussian Smoothed filtered global DAPI channel. Every Tissue Image Region was resized to exclude the very peripheral area of the sections. Sliding parabola filtered DAPI channel within the resized tissue area was used to find nuclei; and Phalloidin was used to identify a cytoplasm (see fig 4B). Mean intensity of MUC1 and Phalloidin was calculated in each cell, and averaged per cell for the entire tissue sample. The average MUC1 signal was then normalized to Phalloidin signal to take in account variability in slide staining conditions.

### High Content Screening

The automated high content screening system consisted of robotic arms; plate stackers; a HighRes Pin Tool; Liconic incubators; Biotek plate washers; dedicated Thermo Fisher Combi Multidrop dispensers for each assay reagent; and PerkinElmer High Content Imaging Instrument Opera Phenix, all choreographed by Cellario software. Cell fixation and immunostaining were all performed in a custom-designed light-protected hood (HighRes Biosolutions). Data analysis and representation was performed using Genedata Screener (Genedata AG) and Spotfire (TIBCO).

For the immunofluorescence screen, P cells were seeded 24 h prior to compound treatment at a density of 12,000 cells/well in 384 well CellCarrier Ultra plates (Perkin Elmer), pre-coated with 0.25 mg/mL Synthemax II SC Substrate (Corning). Compounds of the repurposing library set (Corsello et al., 2017) were used at either 5 doses (35, 3.5, 0.35, 0.035 and 0.0035 μM) or 10 doses (16, 5.6, 1.8, 0.6, 0.21, 0.07, 0.02, 0.008, 0.002 and 0.0008 μM) as indicated. The compounds were transferred in replicate from compound source plates to the cell plates using the HighRes Pin Tool. DMSO was used as a negative control and JQ1 (250 nM) (a bromodomain inhibitor) as positive control, based on earlier studies showing potent reduction of total MUC1 mRNA levels (data not shown). After 48 h incubation, cells were fixed 20 min in 4% PFA (Electron Microscopy Sciences) in PBS, washed twice, then permeabilized (10 min) with 0.5% Triton X-100 (Sigma-Aldrich) in PBS and washed once more. Cells were blocked for 10 min at RT with Blocking solution (100mM Tris HCL pH8; 150mM NaCL; 5g/L Blocking Reagent [Roche]), then incubated 90 min at RT with 1:2000, monoclonal mouse anti-MUC1 (214D4) antibody (Millipore) in Roche Blocking solution, followed by four PBS wash cycles. Then the secondary antibody Alexa Fluor® 546 Goat anti-mouse IgG, Thermo Fisher Scientific and Hoechst 33342 stain, Thermo Fisher Scientific, were applied at a 1:1000 dilution in Roche blocking solution and incubated at RT for 45 min, followed by four PBS wash cycles. Plates were then sealed with a Plate Loc plate and stored in a Liconic incubator at 10°C until imaging.

Image acquisition and analysis was as described in the Fluorescence image acquisition and analysis section. Upon image analysis, two parameters were selected, i) total MUC1 cytoplasmic intensity and ii) cell number as was detected by Hoechst 33342 stained nuclei. MUC1 levels in the presence of DMSO or of JQ1 were defined, respectively as 0 and -100% activity. Values for test compounds were normalized accordingly. Cell number was normalized to DMSO control. All compound concentrations showing > -20% reduction in cell number were masked out. Based on ± 3 median absolute deviation value, hit calling criteria for the initial 5 doses screen were chosen as MUC1 reduction > 30% in 2 or more consecutive concentrations for both replicates. For the initial 10 doses screen, dose response curves were generated for each parameter using Genedata Screener (Genedata AG), and positive hits for the profiling screens were selected based on the compound’s activity in reducing MUC1 abundance without cell toxicity.

For the RT-PCR-based screen (Bittker, 2012), P cells seeded at 2000 cells/well in 384-well, clear bottom, white wall plates were grown for 24 h, then treated with profiling compounds transferred by pinning to duplicate plates. JQ1 (250 nM) and DMSO were used for controls as above. After 24 h, cells were washed and cDNAs generated using ABI Cells-to-Ct kit (Thermo Fisher Scientific, Waltham, MA). MUC1 and HMBS delta Cp values were determined using a Roche LightCycler 480 Instrument in 5µL reactions using TaqMan probes for MUC1 FAM (4351368 assay ID Hs00159357_m1) and HMBS VIC (4448486-assay ID Hs00609297_m1) (Thermo Fisher Scientific). The fold change effect of the compounds on total MUC1 mRNA was normalized to JQ1 and DMSO controls, as described above.

For the viability profiling screen, P cells were seeded 12 h prior to profiling compound treatment at a density of 12,000 cells/well in 384 well Cell Carrier Ultra plates (Perkin Elmer), pre-coated with 0.25 mg/mL Synthemax II SC Substrate (Corning). After 24 h, CellEvent Caspase-3/7 Green Detection Reagent (Thermo Fisher Scientific) and DRAQ7 (Biostatus) were added at 1:5000 final dilution. Cells were imaged daily during 7 days to monitor viability. Image acquisition was done as described below and viability was assessed as number of live cells at the day 6, when most of DMSO treated wells reached about 95% of confluence.

### Mice

Adult, 7-week-old male C57BL/6J mice were purchased from Jackson Laboratory (Bar Harbor, ME) and maintained in specific pathogen-free conditions at the Beth Israel Deaconess Medical Center (BIDMC) and allowed to acclimate for 1 week before use in experiments. All mice used in this study were 8–12 weeks old.

### Administration of SYK inhibitor R788

SYK inhibitor R788 was provided by Rigel Pharmaceuticals (South San Francisco, CA). Mice chow was prepared by Research Diets (New Brunswick, NJ). C57BL/6J mice were fed chow containing 3 g/kg R788 ad libitium for 10 days before experimentation. Control mice were fed normal chow.

### Mesenteric Ischemia-Reperfusion (I/R)

All animal procedures were performed in accordance with the guidelines and approval of the Institutional Animal Care and Use Committee (IACUC) of the BIDMC. Mice were randomly assigned to sham or I/R groups and were anesthetized by intraperitoneal injection of 72 mg/kg pentobarbital. Animals were subjected to I/R, as previously described (Pamuk et al., 2010). Mice were anesthetized with 72 mg/kg nembutal (Butler Schein Animal Health) given i.p. Additionally, 36 mg/kg nembuttal was given s.c. during the experiment as needed to maintain anesthesia. All procedures were performed on anesthetized, spontaneously breathing animals with body temperature maintained at 37°C with a controlled heating pad. A midline laparotomy was performed, and the superior mesenteric artery was identified and isolated. Ischemia was induced by application of ∼85 g of pressure fir 30 min via a small nontraumatic vascular clamp (Roboz Surgical Instruments, Gaithersburg, MD). After 30 min of ischemia, the clamp was removed, the laparotomy incision was repaired with 4-0 Sofsilk (Covidien, Mansfield, MA), the mice were resuscitated with 1.0 ml of prewarmed sterile PBS s.c., and the intestine was allowed to reperfuse for 180 min. Sham-operated mice were subjected to the same operative procedure as the experimental group except that clamping of the superior mesenteric artery was not performed. At the conclusion of the reperfusion period, mice were euthanized by carbon dioxide asphyxiation, following the IACUC Guidelines of the BIDMC. Lung removal consisted of intact extraction of the bronchial tree after expansion with tracheal administration of 200–300 of ice-cold 10% phosphate-buffered formalin and fixed overnight in 10% phosphate-buffered formalin at 4°C. Formalin-fixed lung tissues were washed extensively in PBS, processed, and embedded in paraffin for immunohistochemical analysis.

### Immunohistochemistry

Immunohistochemical staining was performed on formalin-fixed paraffin sections of lung tissues. The samples were subjected to rehydration and antigen retrieval by overnight immersion in 10mM citric acid buffer (pH 6), for overnight at 60° C. Following antigen retrieval, endogenous peroxidase was blocked with 0.3% H2O2 for 15 min followed by blocking with 2.5% fetal bovine serum (FBS) in PBS for 30 min. Sections were then incubated at 4 C overnight with primary antibodies in 2.5% BSA in PBS ((1:500, monoclonal Armenian hamster anti-MUC1, Abcam; 1:400, Rhodamine Phalloidin, Cytoskeleton Inc). Following washing, the sections were incubated for 1 h at 37° C with secondary antibody diluted in 2.5% BSA in PBS (1:500, Alexa Fluor® 488-conjugated AffiniPure Goat anti-Armenian hamster IgG, Jackson Immunoresearch; 1:500). Slides were washed and mounted in ProLong Gold Antifade Mountant with DAPI (Thermo Fischer Scientific).

### Statistical analysis

Statistical analysis was performed and presented using Graphpad Prism version 7.0 software. All data are presented as means ± standard deviation for ‘n’ experiments unless otherwise specified in the figure legends. The exact value of ‘n’ for each experiment can be found in the figure legends. Statistical comparisons of two groups for a single variable with normal distributions were analyzed by unpaired t test. Statistical comparisons of two or more groups with one independent variable were analyzed by One-way ANOVA with Tukey post-tests. Statistical comparisons of two or more groups with two independent variables were analyzed by Two-way ANOVA with Tukey post-tests. *p < 0.05 **p < 0.01 ***p < 0.001 ****p < 0.0001

## References

Argade, A., Bhamidipati, S., Li, H., Sylvain, C., Clough, J., Carroll, D., Keim, H., Braselmann, S., Taylor, V., Zhao, H., et al. (2015). Design, synthesis of diaminopyrimidine inhibitors targeting IgE- and IgG-mediated activation of Fc receptor signaling. Bioorg Med Chem Lett 25, 2122–2128.

Ashburn, T.T., and Thor, K.B. (2004). Drug repositioning: identifying and developing new uses for existing drugs. Nature Reviews Drug Discovery 3, 673–683.

Bittker, J.A. (2012). High-Throughput RT-PCR for small-molecule screening assays. Curr Protoc Chem Biol 4, 49–63.

Bussel, J., Arnold, D.M., Grossbard, E., Mayer, J., Trelinski, J., Homenda, W., Hellmann, A., Windyga, J., Sivcheva, L., Khalafallah, A.A., et al. (2018). Fostamatinib for the treatment of adult persistent and chronic immune thrombocytopenia: Results of two phase 3, randomized, placebo-controlled trials. Am J Hematol 93, 921–930.

Cao, B., Wang, Y., Wen, D., Liu, W., Wang, J., Fan, G., Ruan, L., Song, B., Cai, Y., Wei, M., et al. (2020). A Trial of Lopinavir-Ritonavir in Adults Hospitalized with Severe Covid-19. N Engl J Med.

Castro, M., King, T.S., Kunselman, S.J., Cabana, M.D., Denlinger, L., Holguin, F., Kazani, S.D., Moore, W.C., Moy, J., Sorkness, C.A., et al. (2014). Effect of vitamin D3 on asthma treatment failures in adults with symptomatic asthma and lower vitamin D levels: the VIDA randomized clinical trial. JAMA 311, 2083–2091.

Corsello, S.M., Bittker, J.A., Liu, Z., Gould, J., McCarren, P., Hirschman, J.E., Johnston, S.E., Vrcic, A., Wong, B., Khan, M., et al. (2017). The Drug Repurposing Hub: a next-generation drug library and information resource. Nat Med 23, 405–408.

d’Alessandro, M., Bergantini, L., Cameli, P., Lanzarone, N., Antonietta Mazzei, M., Alonzi, V., Sestini, P., and Bargagli, E. (2020). Serum KL-6 levels in pulmonary Langerhans’ cell histiocytosis. Eur J Clin Invest, e13242.

Dong, E., Du, H., and Gardner, L. (2020). An interactive web-based dashboard to track COVID-19 in real time. Lancet Infect Dis.

Dvela-Levitt, M., Kost-Alimova, M., Emani, M., Kohnert, E., Thompson, R., Sidhom, E.H., Rivadeneira, A., Sahakian, N., Roignot, J., Papagregoriou, G., et al. (2019). Small Molecule Targets TMED9 and Promotes Lysosomal Degradation to Reverse Proteinopathy. Cell 178, 521–535 e523.

Geahlen, R.L. (2014). Getting Syk: spleen tyrosine kinase as a therapeutic target. Trends Pharmacol Sci 35, 414–422.

Grein, J. (2020). Compassionate Use of Remdesivir for Patients with Severe Covid-19. New England Journal of Medicine.

Hilgendorf, I., Eisele, S., Remer, I., Schmitz, J., Zeschky, K., Colberg, C., Stachon, P., Wolf, D., Willecke, F., Buchner, M., et al. (2011). The oral spleen tyrosine kinase inhibitor fostamatinib attenuates inflammation and atherogenesis in low-density lipoprotein receptor-deficient mice. Arterioscler Thromb Vasc Biol 31, 1991–1999.

Ishizaka, A., Matsuda, T., Albertine, K.H., Koh, H., Tasaka, S., Hasegawa, N., Kohno, N., Kotani, T., Morisaki, H., Takeda, J., et al. (2004). Elevation of KL-6, a lung epithelial cell marker, in plasma and epithelial lining fluid in acute respiratory distress syndrome. Am J Physiol Lung Cell Mol Physiol 286, L1088–1094.

Kato, K., Chang, E.H., Chen, Y., Lu, W., Kim, M.M., Niihori, M., Hecker, L., and Kim, K.C. (2020). MUC1 contributes to goblet cell metaplasia and MUC5AC expression in response to cigarette smoke in vivo. Am J Physiol Lung Cell Mol Physiol.

Kato, K., Lillehoj, E.P., Lu, W., and Kim, K.C. (2017). MUC1: The First Respiratory Mucin with an Anti-Inflammatory Function. J Clin Med 6.

Kunwar, S., Devkota, A.R., and Ghimire, D.K. (2016). Fostamatinib, an oral spleen tyrosine kinase inhibitor, in the treatment of rheumatoid arthritis: a meta-analysis of randomized controlled trials. Rheumatol Int 36, 1077–1087.

Li, Y.Y., Hsieh, L.L., Tang, R.P., Liao, S.K., and Yeh, K.Y. (2009). Macrophage-derived interleukin-6 up-regulates MUC1, but down-regulates MUC2 expression in the human colon cancer HT-29 cell line. Cell Immunol 256, 19–26.

McAdoo, S.P., Prendecki, M., Tanna, A., Bhatt, T., Bhangal, G., McDaid, J., Masuda, E.S., Cook, H.T., Tam, F.W.K., and Pusey, C.D. (2020). Spleen tyrosine kinase inhibition is an effective treatment for established vasculitis in a pre-clinical model. Kidney Int.

McAdoo, S.P., Reynolds, J., Bhangal, G., Smith, J., McDaid, J.P., Tanna, A., Jackson, W.D., Masuda, E.S., Cook, H.T., Pusey, C.D., et al. (2014). Spleen tyrosine kinase inhibition attenuates autoantibody production and reverses experimental autoimmune GN. J Am Soc Nephrol 25, 2291–2302.

McAdoo, S.P., and Tam, F.W. (2011). Fostamatinib Disodium. Drugs Future 36, 273.

Mendes, A.I., Matos, P., Moniz, S., Luz, S., Amaral, M.D., Farinha, C.M., and Jordan, P. (2011). Antagonistic regulation of cystic fibrosis transmembrane conductance regulator cell surface expression by protein kinases WNK4 and spleen tyrosine kinase. Mol Cell Biol 31, 4076–4086.

Mocsai, A., Ruland, J., and Tybulewicz, V.L. (2010). The SYK tyrosine kinase: a crucial player in diverse biological functions. Nat Rev Immunol 10, 387–402.

Moore, J.B., and June, C.H. (2020). Cytokine release syndrome in severe COVID-19. Science 368, 473–474.

Muus, C. (2020). Integrated analyses of single-cell atlases reveal age, gender, and smoking status associations with cell type-specific expression of mediators of SARS-CoV-2 viral entry and highlights inflammatory programs in putative target cells. BioRxiv.

Na, H.G., Bae, C.H., Choi, Y.S., Song, S.Y., and Kim, Y.D. (2016). Spleen tyrosine kinase induces MUC5AC expression in human airway epithelial cell. Am J Rhinol Allergy 30, 89–93.

Nakashima, T., Yokoyama, A., Ohnishi, H., Hamada, H., Ishikawa, N., Haruta, Y., Hattori, N., Tanigawa, K., and Kohno, N. (2008). Circulating KL-6/MUC1 as an independent predictor for disseminated intravascular coagulation in acute respiratory distress syndrome. J Intern Med 263, 432–439.

Newland, A., Lee, E.J., McDonald, V., and Bussel, J.B. (2018). Fostamatinib for persistent/chronic adult immune thrombocytopenia. Immunotherapy 10, 9–25.

Ohtsuki, Y., Fujita, J., Hachisuka, Y., Uomoto, M., Okada, Y., Yoshinouchi, T., Lee, G.H., Furihata, M., and Kohno, N. (2007). Immunohistochemical and immunoelectron microscopic studies of the localization of KL-6 and epithelial membrane antigen (EMA) in presumably normal pulmonary tissue and in interstitial pneumonia. Med Mol Morphol 40, 198–202.

Pamuk, O.N., Lapchak, P.H., Rani, P., Pine, P., Dalle Lucca, J.J., and Tsokos, G.C. (2010). Spleen tyrosine kinase inhibition prevents tissue damage after ischemia-reperfusion. Am J Physiol Gastrointest Liver Physiol 299, G391–399.

Pushpakom, S., Iorio, F., Eyers, P.A., Escott, K.J., Hopper, S., Wells, A., Doig, A., Guilliams, T., Latimer, J., McNamee, C., et al. (2019). Drug repurposing: progress, challenges and recommendations. Nat Rev Drug Discov 18, 41–58.

Ruan, Q., Yang, K., Wang, W., Jiang, L., and Song, J. (2020). Clinical predictors of mortality due to COVID-19 based on an analysis of data of 150 patients from Wuhan, China. Intensive Care Med.

Sakai, M., Kubota, T., Ohnishi, H., and Yokoyama, A. (2013). A novel lung injury animal model using KL-6-measurable human MUC1-expressing mice. Biochem Biophys Res Commun 432, 460–465.

Singh, R., Masuda, E.S., and Payan, D.G. (2012). Discovery and Development of Spleen Tyrosine Kinase (SYK) Inhibitors. Journal of Medicinal Chemistry 55, 3614–3643.

Smith, J., McDaid, J.P., Bhangal, G., Chawanasuntorapoj, R., Masuda, E.S., Cook, H.T., Pusey, C.D., and Tam, F.W. (2010). A spleen tyrosine kinase inhibitor reduces the severity of established glomerulonephritis. J Am Soc Nephrol 21, 231–236.

Uhlen, M., Fagerberg, L., Hallstrom, B.M., Lindskog, C., Oksvold, P., Mardinoglu, A., Sivertsson, A., Kampf, C., Sjostedt, E., Asplund, A., et al. (2015). Proteomics. Tissue-based map of the human proteome. Science 347, 1260419.

Vestbo, J. (2002). Epidemiological studies in mucus hypersecretion. Novartis Found Symp 248, 3-12; discussion 12-19, 277-282.

Weinblatt, M.E., Kavanaugh, A., Genovese, M.C., Musser, T.K., Grossbard, E.B., and Magilavy, D.B. (2010). An Oral Spleen Tyrosine Kinase (Syk) Inhibitor for Rheumatoid Arthritis. New England Journal of Medicine 363, 1303–1312.

Wilson, J.G. (2020). Cytokine profile in plasma of severe COVID-19 does not differ from ARDS and sepsis. medRxiv.

Yi, Y.S., Son, Y.J., Ryou, C., Sung, G.H., Kim, J.H., and Cho, J.Y. (2014). Functional roles of Syk in macrophage-mediated inflammatory responses. Mediators Inflamm 2014, 270302.

Zhang, C., Wu, Z., Li, J.W., Zhao, H., and Wang, G.Q. (2020). The cytokine release syndrome (CRS) of severe COVID-19 and Interleukin-6 receptor (IL-6R) antagonist Tocilizumab may be the key to reduce the mortality. Int J Antimicrob Agents, 105954.

Zhou, F., Yu, T., Du, R., Fan, G., Liu, Y., Liu, Z., Xiang, J., Wang, Y., Song, B., Gu, X., et al. (2020). Clinical course and risk factors for mortality of adult inpatients with COVID-19 in Wuhan, China: a retrospective cohort study. Lancet 395, 1054–1062.

